# CLARITY-compatible lipophilic dyes for electrode marking and neuronal tracing

**DOI:** 10.1101/061135

**Authors:** Kristian H. R. Jensen, Rune W. Berg

## Abstract

Fluorescent lipophilic dyes, such as DiI, stain cellular membranes and are used extensively for retrograde/anterograde labelling of neurons as well as for marking the position of extracellular electrodes after electrophysiology. Convenient histological clearing techniques, such as CLARITY, enable immunostaining and imaging of large volumes for 3D-reconstruction. However, such clearing works by removing lipids and, as an unintended consequence, also removes lipophilic dyes. To remedy this wash-out, the molecular structure of the dye can be altered to adhere to both membranes and proteins so the dye remains in the tissue after lipid-clearing. Nevertheless, the capacity of such modified dyes to remain in tissue has not yet been tested. Here, we test dyes with molecular modifications that make them aldehyde-fixable to proteins. We use three Dil-analogue dyes, CM-DiI, SP-DiI and FM 1-43FX that are modified to be CLARITY-compatible candidates. We use the challenging adult, myelin-rich spinal cord tissue, which requires prolonged lipid-clearing, of rats and mice. All three dyes remained in the tissue after lipid-clearing, but CM-DiI had the sharpest and FM 1-43FX the strongest fluorescent signal.

## Introduction

Tracing the position of an extracellular electrode after electrophysiological experiments in the nervous system is important to verify both the brain region and the distance between electrode and nearby neurons, which are often identified via either genetically expressed reporters or immunohistochemistry. The tracing is commonly accomplished using a non-toxic fluorescent and lipophilic dye, as the widely used carbocyanine dye DiI, which is painted on the electrode shank prior to insertion. The electrode then leaves remnants of the dye that can be easily be detected by fluorescent microscopy in histological sections.^1–3^ An additional widespread application of lipophilic fluorescent dyes is for tracing neuronal connections and projections in the nervous system^4,5^especially since these dyes, e.g. DiI, are well-suited for immunohistochemistry^6^ and have minimal photo-bleaching.^7^ DiI can be transported retrogradely^8^ and has been used for post-mortem axonal labelling in animals^9^ and humans.^10,11^Nevertheless, this approach has an unfortunate limitation due to the need to do serial sectioning of the tissue, which is imprecise and labour intensive. The tracing can require several sections if the cutting angle is not parallel to the electrode track and includes the risk of distorting the samples.

However, recent developed clearing techniques, such as CLARITY,^12^ PACT,^13^ CUBIC,^14^ iDISCO,^15^ and Sca/e^16^ conveniently allow immunohistochemistry, imaging and 3D reconstruction in large volumes of unsectioned tissue.^12,17,18^These clearing techniques work essentially by washing away the lipids with detergents and solvents. CLARITY works by polymerizing the fixed tissue into an acrylamide hydrogel prior to lipid-removal. This leaves molecules with amine ends, i.e. proteins, DNA, and RNA, into a protein skeleton structure. These lipid-clearing processes introduce a caveat when combined with the axonal tracing or electrode marking: The lipophilic dyes are also washed out since they adhere to the lipids.^19–21^CLARITY and similar clearing techniques are thus currently incompatible with the employment of lipophilic dyes in tracing and marking. Clearing alternatives such as *Clear*^T 17,22^do not wash out the lipids and therefore leaves the lipophilic dyes secured, but they do not clear as well as CLARITY and therefore have low visualization depth. The clearing methods SeeDB,^23^ 3DISCO^24^ and BABB^25^ have high visualization depth, but the penetration depth of immunohistochemistry is limited to 100–250 *μ*m^26^ or lower.^18^ It would therefore be an advantage to find alternative dyes for tracing, which would be compatible with CLARITY and similar clearing techniques^15^ while also having a strong tie to the cellular membranes to limit the diffusion before fixing. Notably, a very recent improvement of the Sca/eA clearing technique, called Sca/eS, is able to overcome these shortcomings and preserve DiI-labeling.^21^ This was demonstrated in axon tracks, which were labelled with DiI-crystals applied onto the tissue surface on *post-mortem* brain tissue. The tissue was fixed prior to the dye-application and then fixed again for a week.^21^ In spite of these improvements, fixability of lipophilic dyes, that remain in the tissue after lipid-clearing, continue to be a desirable quality.

The simplest approach to obtain an alternative dye is to chemically change an existing dye, e.g. DiI, into a DiI—analogue, which possesses both the property of adhering to the cellular membranes and protein structures, such that it would remain in the tissue after lipid-clearing. Such dyes have already been developed, in particular the analogues of DiI: CM-DiI,^20,27^ SP-DiI^28^ and FM 1-43FX,^29–31^with modifications that make them aldehyde-fixable to proteins (Fig. 1). Here, we tested the DiI-analogues on multi-electrodes placed in the spinal cord^32^ using CLARITY as clearing method. The spinal cord has an envelope of dense white matter and is therefore a difficult tissue to clear of lipids, though it has successfully been cleared previously.^33^ We also imaged and 3D-reconstructed the spinal cord and dye traces using confocal microscopy.

**Figure 1.**
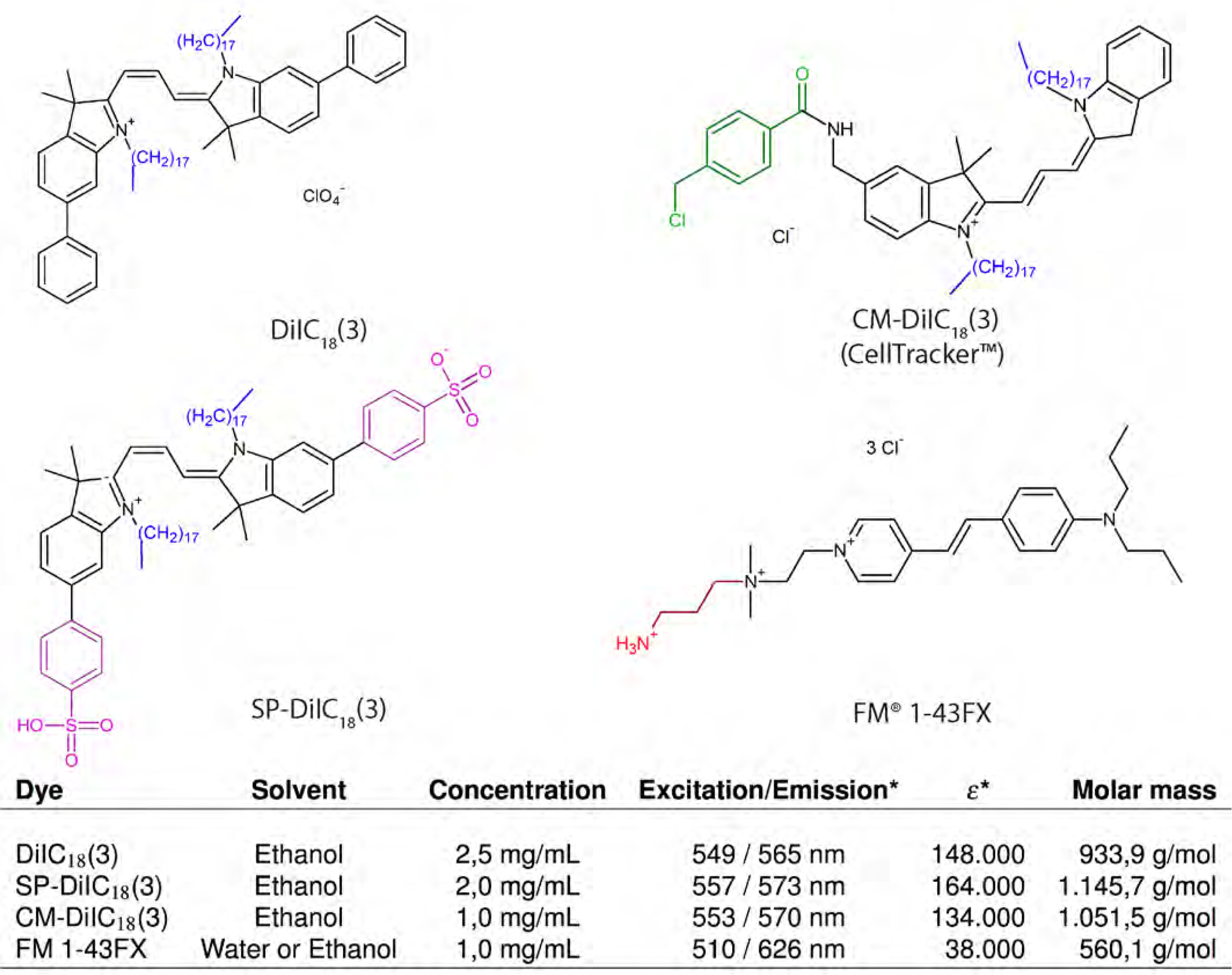
Overview of chemical structures and properties of DiI and CLARITY-compatible analogues. DiIC_18_(3) (DiI in short) is a fluorescent lipophilic cationic indocarbocyanine dye used for single molecule imaging, fate mapping, and neuronal tracing, as it is retained in the lipid bilayers. CM-DiIC_18_(3) (CM-DiI in short) incorporates a mildly thiol-reactive chloromethylbenzamido (CM) substituent (green) that confers aldehyde fixability via conjugation to thiol-containing proteins. SP-DiIC_18_(3) (SP-DiI in short) has two sulfophenyl (SP) groups (magenta) that confers fixability and a greater ε than DiIC_18_(3). FM 1-43FX, which is less similar to DiIC_18_(3), is a lipophilic styryl dye with an aliphatic amine side chain (red) for aldehyde fixability. It has, notably the lowest ε, and the greatest Stokes shift, but also half the molar mass due to lacking the two 18-carbon-long alkyl tails (blue). *Spectral properties determined in methanol. ε: molar attenuation coefficient in *cm*^-1^*M*^-1^.

## Results

### Fixable lipophilic dyes remain after lipid-clearing

We tested three dye-candidates, the dyes CM-DiI, FM 1-43FX, and SP-DiI (Fig. 1), in living tissue. Multi-electrodes, which had been dipped in the dye (diluted in ethanol), were inserted for 10 min into the spinal cord of an anesthetized mouse (Fig. 2a-c). Traces of the fixable dyes remained in the tissue as indicated by the fluorescent signals (Fig. 2d-e) after lipid-clearing of the spinal cord using CLARITY (Fig. 2f-g). In the widefield microscope, dye FM 1-43FX had the broadest and most vivid overall fluorescence, while CM-DiI had the sharpest traces. SP-DiI was barely visible with a poor overall fluorescent signal (Fig. 2d-e). Twelve traces were identifiable for CM-DiI, 13 for FM 1-43FX, and only 9 for SP-DiI, where 14 were expected for each dye. Electrode track marking using CM-DiI and FM 1-43FX were the most reliable (Fig. 3). They both enabled clear electrode localization in 3D (Fig. 3d,h), whereas the SP-DiI reconstruction required additional post-processing when imaged together with DAPI (subtraction of the spectrally overlapping DAPI signal, Fig. 3j-l) and was limited by the low fluorescence and few identifiable traces (Fig. 3i-l). For the electrodes with CM-DiI, 86% of the tracks were recovered, and 93% of the tracks for the electrodes applied FM 1-43FX (Fig. 4a).

In euthanized rats, where the spinal cord had been PFA-fixed prior to insertion of electrodes for 30 min, there was a strong SP-DiI fluorescence after CLARITY-clearing (Supplementary Fig. 1).

**Figure 2.**
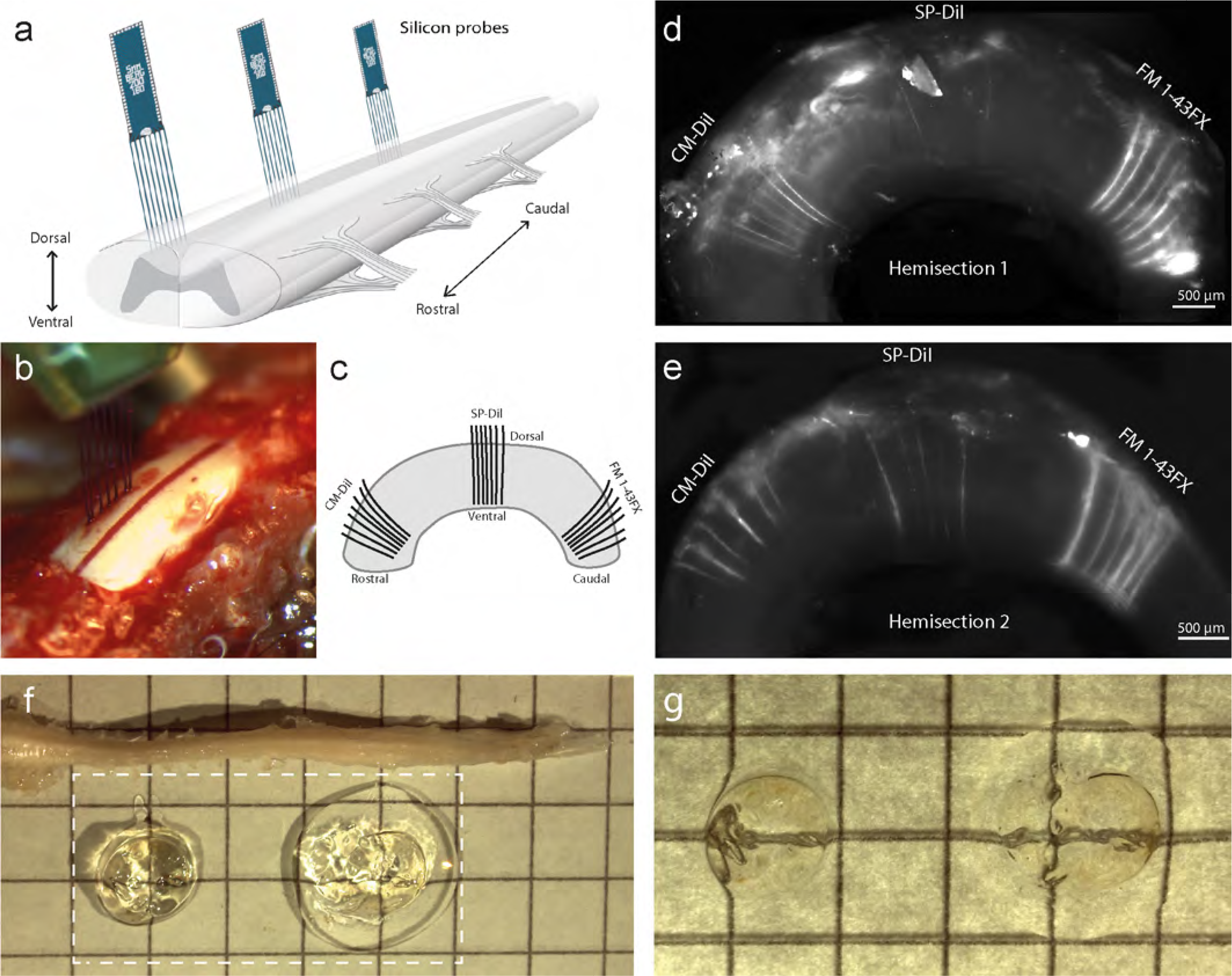
DiI-analogues remained in CLARITY-treated nervous tissue and reveal electrode placement. (a) Multielectrodes painted with either CM-DiI, SP-DiI or FM 1-43FX are inserted from the dorsal side in a sagittal plane into the spinal cord for 10 minutes each. (b) The spine is surgically opened to expose the lumbar spinal cord in the anaesthetized mouse with one electrode array inserted. (c) Illustration of the location of the electrodes tracks covered with different dyes from a sagittal view. (d-e) Widefield images of the hemicords demonstrate that fluorescent DiI-analogues remain in the tissue after CLARITY treatment. FM 1-43FX (right) has the strongest overall fluorescent signal, and CM-DiI (left) has the sharpest traces. (f) Images of the CLARITY cleared hemisections (broken box) imaged in a drop of 80% glycerol on a glass slide with PFA-fixed spinal cord above for comparison. The hemisections unavoidably curl up into circles. (g) Close up of hemisections imaged with backlighting. The multielectrode had seven shanks, each 50 *μ*m thick and 200 *μ*m apart.

**Figure 3.**
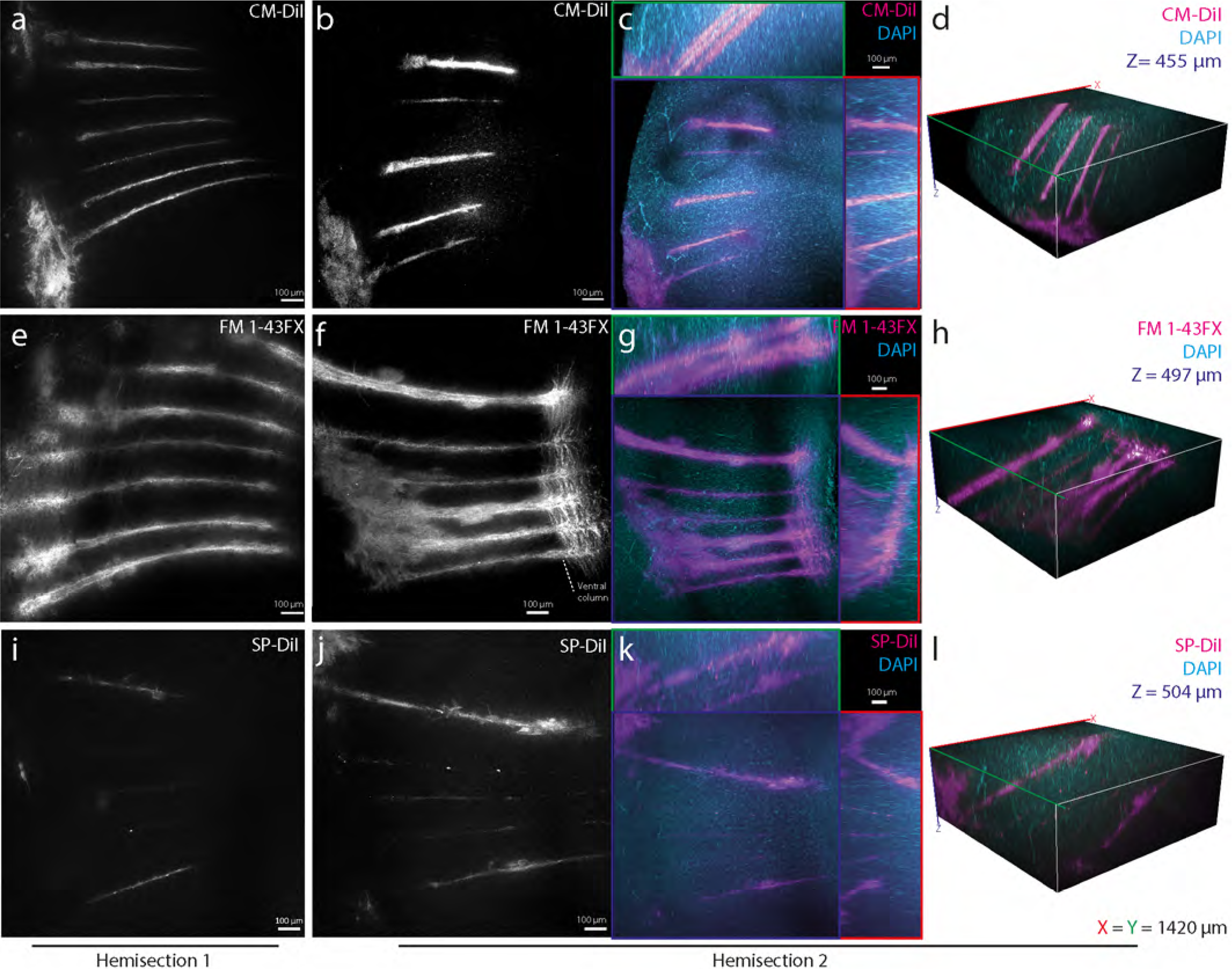
Comparing CLARITY-compatible lipophilic dyes for tracing and 3D-visualization. Confocal images of CLARITY-processed mouse spinal cord (a, b) show clear traces of CM-DiI and (c, d) with both CM-DiI (magenta) and DAPI (cyan) in Z(blue), X(red) and Y(green) planes and 3D-reconstructions (d). (e-h) FM 1-43FX staining, which also stains the white matter column in longitudinal direction at the electrode tip (right, f). Both CM-DiI and FM 1-43FX enable clear electrode localization in 3D (d, h). (i-l) SP-DiI had faint traces, that were difficult to distinguish from the background and localize in 3D (l). All images positioned with dorsal surface left, and images are from hemisection 2 (except a, e, i). Diffuse staining to the left is the superficial tissue at the point of entry. The multielectrode had seven shanks, each 50 Um thick and 200 Um apart. The laser power used to excite CM-DiI and FM 1-43FX was 1.25 mW and 2.5 mW for SP-DiI. Due to the high excitation laser power, the broad spectra of DAPI and the faint signal of SP-DiI, it was necessary to removed DAPI background fluorescence by subtracting 50% of the DAPI images from the SP-DiI images (j-l). The depths of the imaging plane from the surface are given as z-stack midpoints (hemisection 1 and 2): 915 Um and 805 Um (CM-DiI), 910 Um and 706 Um (SP-DiI), 667 Um and 675 Um (FM 1-43FX). (a: Maximum Intensity Projection of 53 images, z= 338 Um, b-d: 66 images, z=455 Um, e: 53 images, z=335 Um, f-h: 72 images, z=497 Um, i: 31 images, z=200 Um, j-l: 73 images, z=504 Um)

**Figure 4.**
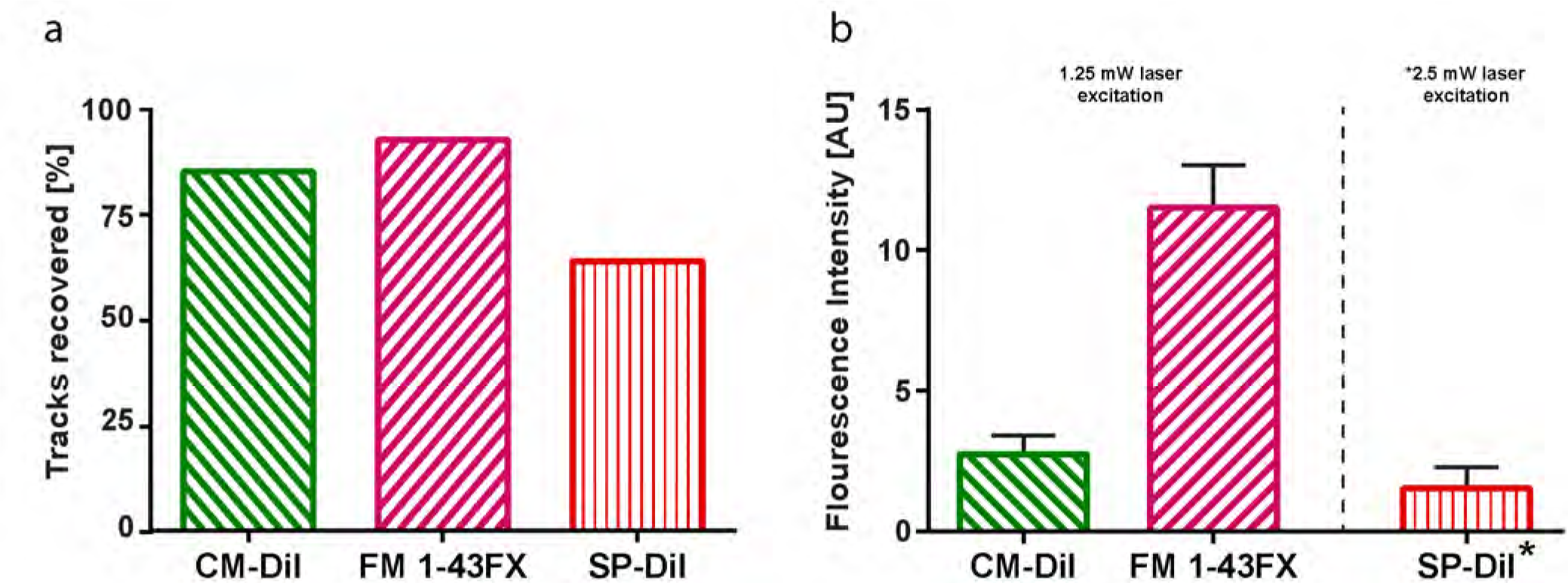
Performance of the DiI-analogues quantified by the fluorescent residue from the electrode tracks after lipid-clearing. (a) Electrode tracks recovered by visual inspection from the two hemisections: twelve traces were identifiable for CM-DiI (86%), 13 for FM 1-43FX (93%), and only nine for SP-DiI (93%) out of 14. (b) Fluorescence intensity in rectangles placed on the identified electrode traces with background fluorescence subtracted. Values are the mean intensity of the identified tracks with the standard error of the mean. *The laser power used to excite SP-DiI was double (2.5 mW) that of CM-DiI and FM 1-43FX (1.25 mW) and therefore is not directly comparable, hence the broken line of separation.

As a control, we tested whether a non-fixable DiI variant, DID, the redshifted version of DiI, remained in the tissue following lipid-clearing using both CLARITY and Sca/eA2. DiD was applied on electrodes prior to insertion into the tissue. The tissue was then PFA-fixed and after lipid-clearing using both CLARITY (Supplementary Fig. 2) and Sca/eA2 (data not shown) there was no traceable remnants of the dye in the tissues.

### FM 1-43FX has the strongest fluorescence

Fluorescence measurements demonstrated FM 1-43FX to have the strongest fluorescence signal, followed by CM-DiI (Fig. 3). The fluorescent signals were quantified as the intensity of signal in a rectangle place over each identifiable electrode vestige. The mean signal of these rectangles ± standard error of the mean show a clear difference between the intensities of the three dyes (Fig. 4b). SP-DiI had limited staining and required double the excitation laser power for all images (2.5 mW versus 1.25 mW at 514 nm). Both CM-DiI and FM 1-43FX had stronger fluorescent signals and the electrode traces were visible in imaging depths down to 866 *μ*m (Supplementary Fig. 3). FM 1-43FX has a broader emission spectra than CM-DiI and SP-DiI, which led to crosstalk (bleed-through) when acquiring a multi-channel image with DAPI and GFP. This bleed-through required image post-processing to subtract the separate fluorescent signals (Supplementary Fig. 3). In addition, in order to verify whether using water versus ethanol as the dye solvent changed the degree of dye binding to the electrode and tissue staining we tested the degree of staining using both these dye solvents for FM 1-43FX. The two solvents had no clear difference in the degree of staining (Supplementary Fig. 3). We also tested whether using organic solvents, such as glycerol and 2,2’-thiodiethanol (TDE) as the refractive index matching medium (RIMM) had an effect on the dyes. We compared glycerol and TDE with the aqueous 88% w/v Histodenz in PBS and found no noticeable difference in fluorescence and image quality (data not shown).

### Dye diffusion and axon tracing

The width of the fluorescent electrode traces varied from 5 to 90 *μ*m. CM-DiI and SP-DiI had thinner (<30 *μ*m) traces, while FM 1-43FX had broader traces (>30 *μ*m) (Fig. 5a-c). The broader staining using FM 1-43FX is likely due to the lateral diffusion along the membrane associated with labelling of cells and projections (Fig. 5c). FM 1-43FX extensively stained the white matter column in longitudinal direction at the electrode tip, and single axon projections are clear and possible to delineate (Fig. 5d,e). Both CM-DiI and SP-DiI were constricted to a smaller region around the electrode vestige. This is likely due to the larger molecular weight of both dyes (Fig. 1) combined with the short exposure time. As a result, the dye diffusion was rather limited, even though carbocyanine dyes are known to be transported in axons.^7–11^Nevertheless, the fact that all three dyes remained in the tissue after lipid-clearing combined with the well-established tracing properties of carbocyanine dyes suggest that these dyes would work well for axon tracing and large volume microscopy of tissue processed using CLARITY and similar protocols.

**Figure 5.**
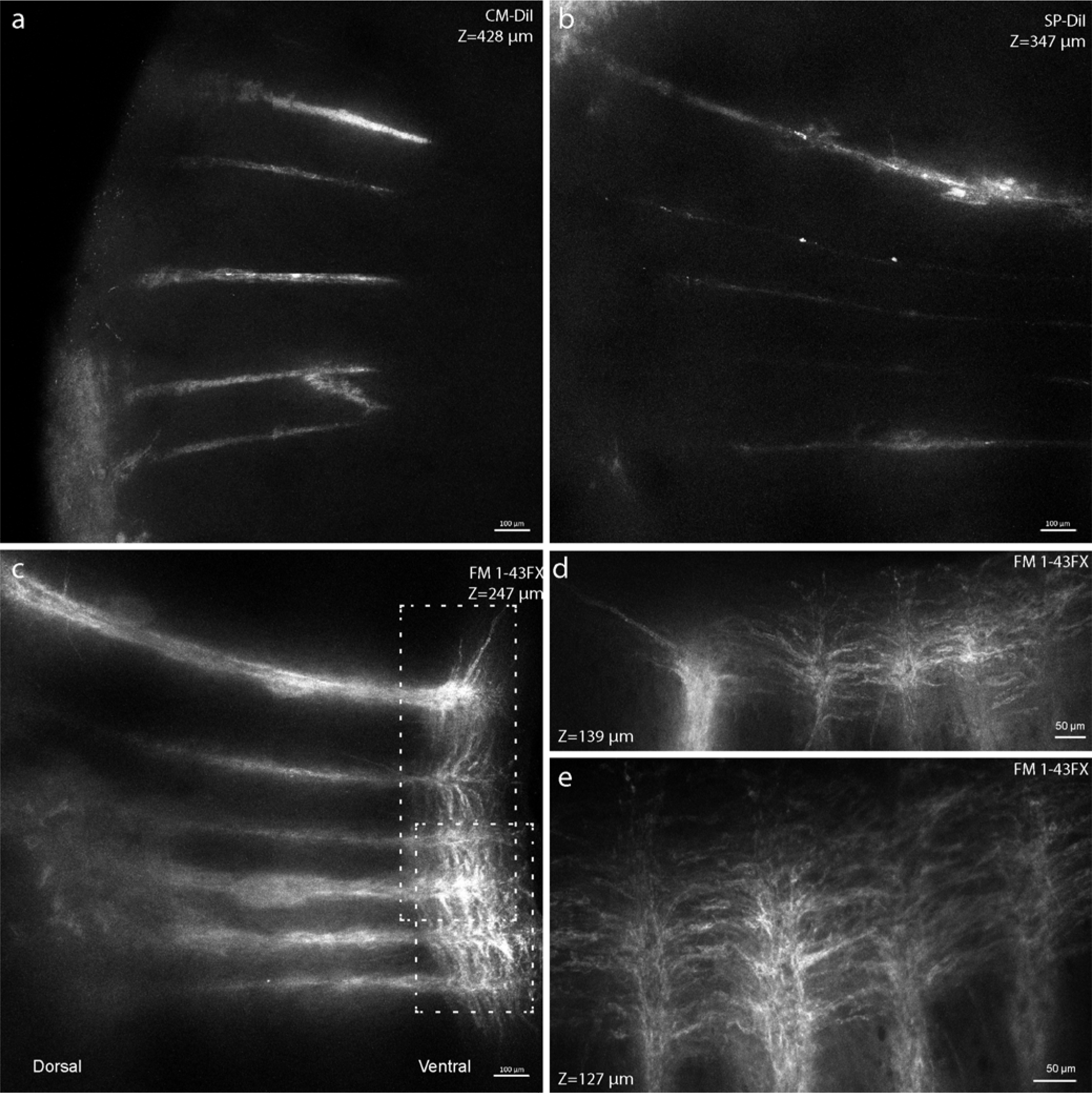
CM-DiI gives sharp staining, while FM 1-43FX gives brighter traces of electrodes and axons. High resolution images of CM-DiI, FM 1-43FX and SP-DiI (as seen in Fig. 3, b, f, j) reveal (a) CM-DiI has the sharpest traces while (b) FM 1-43FX has the strongest overall fluorescence (excitation laser power was 1.25 mW in both). (c) SP-DiI had a fainter signal even with the double laser power (2.5 mW). (d, e) Close-ups from (b, boxes) show the lateral diffusion of FM 1-43FX along the axonal membranes in the ventral white-matter column at the electrode tip. (a: Maximum Intensity Projection of 65 images, z=428 **μ**m, b: 38 images, =247 **μ**m, c: 53 images, z=347 **μ**m, d: 11 images, z=139 **μ**m, e: 10 images, z=127 **μ**m)

## Discussion

It has previously been possible to achieve only two out of three important qualities in histology: High visualization depth, compatibility with immunohistochemistry, or compatibility with lipophilic dyes. Fortunately, with fixable DiI-analogues that are also compatible with lipid-clearing protocols, such as CLARITY, it is now possible to get all three desired qualities. In the present study, we therefore tested the compatibility of three groups of lipophilic dyes with CLARITY, for the purpose of visualizing electrode placement and neuronal projections. Since other lipid-clearing methods work in similar yet gentler ways compared with CLARITY, these fixable DiI-analogues are likely to work with other lipid-clearing methods as well. We traced electrode tracks following electrophysiology and 3D reconstruction with the option of including immunostaining. All three dyes had preserved fluorescent signal even after prolonged lipid-clearing using CLARITY. CM-DiI had the sharpest and FM 1-43FX had the strongest signal (Figs. 2–5), which demonstrates a graceful compatibility with CLARITY. Together our results show that the DiI-analogues, including FM 1-43FX, are fixable, but show different staining patterns and fluorescent intensities (Fig. 3–5). We recommend using FM 1-43FX to achieve the strongest signal and CM-DiI for the sharpest signal.

There are alternative variants of the DiI and FM dyes, which have alterations in their chemical structures. Since the alterations are minor, these alternative dyes likely have similar properties regarding the fixation and lipid-extraction protection. The three lipophilic dyes that were tested in the present report, are representatives of the main types of lipophilic and fixable dyes: 1) the sulfonated DiI-variants, 2) DiI with a chloromethyl benzamide modification and 3) FM dyes with an attached aliphatic amine (Supplementary Fig. 4). These three dyes are suitable to compare since they have similar excitation and emission spectra and could thus be excited and imaged with the same microscope parameters, i.e. excitation laser and emission filter wavelengths. Other variants may have different excitation/emission spectra, but they are likely to have similar chemical properties regarding lipid-clearing, since their fixable parts are the same. CM-DiI and SP-DiI are variants of the commonly used long-chain lipophilic carbocyanine dye DiI. We selected SP-DiI as a representative for the group of anionic sulfonated DiI-variants with sulfo-, DiI-DS, DiD-DS, and sulfophenyl-modifications, SP-DiI, SP-DiO (Supplementary Fig. 4).

An example of a subtle difference between variants is the DiI with different length of the carbon bridge between the indoline rings, which gives different excitation and emission wavelengths e.g. DiD (DiIC_18_(5), ex/em 650/670 nm) a red-shifted version with a 5 rather than 3 carbon bridge (Supplementary Fig. 4). This neither effects fixability, nor does it make the dye less lipophilic. The lipophilic property is dependent on the length of the alkyl chain length (Fig. 1). Similarly, FM 1-43 has a red-shifted variant, FM 4-64, with a 6 rather than 2 carbon bridge between the aromatic rings. Nevertheless, the parts of the molecules, which confer the fixability of the dyes, e.g., the aliphatic amine, remain the same, and therefore the property of remaining in the tissue after lipid-clearing is likely to be conserved. The chloromethyl benzamide (CM) and sulfophenyl (SP) modifications (Fig. 1), keep the dyes retained throughout PFA-fixing and permeabilization procedures with detergents and solvents e.g. Triton-X and acetone.^20^ The mildly thiol-reactive CM substitution confers aldehyde fixability via conjugation to thiol-containing proteins. The mechanism of SP-DiI retention is currently unknown.

SP-DiI has the highest molar attenuation coefficient (Fig. 1) of the lipophilic fluorescent dyes that had tolerated fixing and permeabilization according to the manufacturer (Thermo Fisher Scientific). Nevertheless, SP-DiI only gave faint staining and the electrode tracks were difficult to identify (Figs. 3–5). In addition, SP-DiI required double the excitation laser power compared to CM-DiI and FM 1-43FX during imaging. When we inserted the electrode for 30 minutes in a pre-fixed rat spinal cord, a bright and sharp staining was found (Supplementary Fig. 1), which suggests that SP-DiI staining requires a longer exposure in the tissue. It may, therefore, work better for longer exposure such as during chronic electrode implantations. DiI is commonly used for tracing neuronal tracts in human post-mortem brain samples, and therefore SP-DiI would be a suitable analogue for similar neuronal tract tracing, where lipid-clearing and large volume microscopy was required.

CM-DiI gave a bright and sharp staining of electrodes tracks after only 10 minutes in the tissue (Fig. 2–4). It had limited diffusion into surrounding cell membranes and gave narrow electrode traces. For electrode tracing, CM-DiI was bright, sharp and reliable (Fig. 3a-d) with 12 of the 14 electrode traces recovered (Fig. 3a). However, we observed an overall diffuse staining of CM-DiI when the insertion site was close to the incision of the pia mater (Supplementary Fig. 3). In conclusion, CM-DiI gave a bright and sharp staining, and is therefore well suited as a marker of acute electrode placement, and likely also chronic.

FM 1-43FX also gave a bright staining, despite its low molar attenuation coefficient (38,000 *m^2^/mol*). The molar mass of FM dyes is half of DiI-variants (Fig. 1) and therefore the molar concentration is approximately double compared to CM-DiI in our experiments, which contributes to the stronger fluorescence. FM 1-43FX gave a broader and more diffuse staining of the electrode tracks, as well as lateral diffusion along the white matter fibers (Fig. 2f). Its small molecular weight gives FM 1-43FX the fastest diffusion of three dyes, which in parts could explain the broader staining of the electrode track. It should be noted that the diffusion in the extracellular space is rather limited and anomalous since the molecules are lipophilic and therefore have ’sticky’ boundary conditions of the cellular membranes.^34^ FM dyes are styryl lipophilic dyes used to image synaptic vesicle exocytosis and endocytosis and has an analogous chemical structure to DiI, but notably lacks the heavy 18 carbon alkyl chains. FM 1-43FX is a modified variant of the dye FM 1-43, with a aldehyde-fixable aliphatic amine terminal (Fig. 1). We selected FM 1-43FX since it matched the spectra of DiI and since FM 1-43 is the most used of the FM dyes.^35^ It is worth noting, that FM 1-43, the non-fixable version of FM 1-43FX, is not inert, but can bind to and block muscarinic acetylcholine receptors. Other FM dyes can block the nicotinic muscarinic acetylcholine receptor and pass through and block mechanotransduction channels in the hair cells of the cochlea.^35^

An important property of FM dyes is their rapid lateral diffusion and involvement in vesicle endocytosis. These dyes are absorbed into the outer lipid bilayer and then taken up by endocytosis at the pre-synaptic terminals, especially in active neurons.^35^ The low molar mass and the involvement in endocytosis likely contributes to the spread of staining compared to CM- and SP-DiI (Figs. 1–2). Therefore, when chronically implanting electrodes marked with FM 1-43FX, the dye will likely rapidly stain the neuronal projections and synaptic terminals in addition to the cells at the electrode site. The rapid lateral membrane diffusion may also contribute to the high fluorescence yield by preventing dye aggregation, which is a common cause of fluorescence self-quenching.^36^

DiI dyes are also effective neuronal tracers, but are known to have slow diffusion and therefore require longer incubation.^37^ Their diffusion speed is not specifically dependent on alkyl chain length,^38^ but it can be increased by modifying the alkyl chains to unsaturated alkenes as with *FASTDiI* (DiIΔ^9,12^−C_18_(3)).^37,39^Alternatively, the application of an electrical field or the depletion of cholesterol using e.g. methyl-β-cyclodextrin in the tissue improve lateral diffusion speed.^40^ However, fixable DiI variants are currently not commercially available with alkene chains. It is therefore unlikely that the CM and SP modifications effect diffusion speed and they are thus expected to diffuse and behave like regular DiI.^41,42^Therefore, SP-DiI and CM-DiI would be well suited for chronic electrode implantations.

FM and DiI dyes are considered to have limited cytotoxic and not to inhibit cell proliferation.^3,7^However, one study finds that carbocyanine dyes with long (≥C16) alkyl side chains, as DiI, are toxic to embryonic rat motoneurons. Another study that they inhibit mitochondrial electron transport chain activity.^43^ Therefore, carbocyanine dyes with shorter (≤C12) alkyl side chains may be more suitable for embryonic tissue or long-term labelling. CM-DiI is reportedly toxic to preadipocytes.^44^

There are other clearing methods similar to CLARITY that use organic solvents or detergents, which disrupts membranes and impairs reliable signal preservation of lipophilic dyes i.e. iDISCO,^15^ CUBIC,^14^ Sca/e^16^ and TDE.^45–47^Sca/e uses a mixture of urea and glycerol, which disrupts the membranes and dissolves lipophilic dyes.^16,17^A promising novel type of sorbitol-based clearing method called Sca/eS has been demonstrated to preserve membrane structure.^21^ Remarkably, Sca/eS also preserves DiI-labeling in of axon tracks labelled with DiI-crystals applied onto the tissue, though it should be noted that it was performed on *post-mortem* brain tissue, which was fixed prior to the dye-application, after which the tissue again was fixed a week.^21^ Other alternatives do not damage the membranes or remove lipophilic dyes. For instance, FocusClear^TM^ and *Clear^T^* are based on aqueous solutions that are compatible with lipophilic dyes and immunohistochemistry, but they do not clear as well as CLARITY and Sca/e.^17,22^SeeDB and FRUIT use fructose, which preserves membranes and lipophilic dyes, but the processed tissues has limited permeable for macromolecules and antibodies and are thus incompatible with large volume immunohistochemistry.^19,26,48^

We used the passive CLARITY technique, PACT,^12^ which relies on incubation and agitation at 37–40°C in 8% SDS. We consider the high detergent concentration and heating during PACT to one of the harshest clearing methods. Active CLARITY with electrophoresis produces oxygen radicals making it even harsher than passive CLARITY. These free radicals can chemically reduce the dye and quench fluorescence. However, fluorescent proteins are well preserved in active CLARITY and ACT-PRESTO.^12,49^Since the DiI-analogues are covalently bound to primary amines on proteins and nucleic acids by methylene bridges following aldehyde fixation, they are not removed during lipid extraction by solvents or detergents. The fixable DiI-analogues were stable in CLARITY and the organic solvents i.e. 80% Glycerol and 65% TDE^45^ used as imaging mediums. When stored in 80% glycerol for 6 months the tissue and traces retained their image worthy fluorescent signal and did not require additional laser power to create identical images as presented here. We therefore predict that the fixable DiI-analogues tested in passive CLARITY are also compatible with other clearing methods based on organic solvents or lipid removal as Sca/e,^16^ iDISCO,^15^ CUBIC^14^ and TDE.^45–47^Moreover, these dyes may also be useful in other animal or plant tissues; that require CLARITY clearing with electrophoresis or the addition of enzyme digestions.^50,51^

In general, dyes, as DiI, are used for antero-and retrograde tracing of neurons in vivo, in vitro, and post-mortem to better understand neuronal circuits.^7^ Other common neuronal tracing methods rely on e.g. viral vectors, dextran-conjugates, Cholera Toxin Subunit B or biocytin.^52^ To date, the only neuronal tracer studies in CLARITY have been done using viral vectors.^53,54^ Viral vectors are powerful tools for studying neuronal circuits.^55^ Fixable DiI-analogues could serve as a viable alternative to tracing using viral vectors (See Supplementary Table 1 for an overview of dyes).

## Methods

All experiments and procedures were approved by the Department of Experimental Medicine, and according to procedures laid out by Danish Ministry of Justice and the Danish National Committee for Ethics in Animal Research. Adult Sprague-Dawley rats of both sexes were used. Transgenic mice (PV-Cre:channelrhodopsin-2/EYFP were provided by the Jackson lab via the
Barkat lab.^56^ Mice expressing enhanced green fluorescent protein (eGFP) under the promoter of glial fibrillary acidic protein [GFAP; Tgn(hgFAPEGEP)] GFEC 335) were provided by the Perrier lab^57^ who got the strain from the Kirchhoff lab.^58^

### Tracer Dyes

The multi-electrode was submerged in 0.5 ml of a dye solution in a 1.5 ml microcentrifuge tube (Eppendorf) and the electrode air-dried out over at least 5 min before usage and protected from light under an aluminum cover. The dye solutions were either 0.25% w/v DiIC_18_(3), 0.25% w/v DiDC_18_(5), 0.2% w/v SP-DiIC_18_(3), or 0.1% w/v CM-DiIC_18_(3) (CellTracker^TM^) in ethanol, or 0.1% w/v FM^®^ 1-43FX (a fixable version of FM^®^ 1–43) in ethanol or water (Thermo Fisher Scientific). The electrodes were washed by leaving them overnight in ethanol. When painting onto the electrodes, ethanol is preferable as solvent due to quicker drying of the dye after application. Chemical structures draw with ChemSketch (ACD/Labs).

### Electrode insertion

The multi-electrode (A4x4-3mm-50-125, Neuronexus Inc.) was attached to a micromanipulator and gradually inserted dorsally 0. 3–1 mm into the spinal cords in the sagittal plane slightly laterally from the hemicord midline. In the mouse brain, the multielectrode was inserted approximately 0.2 mm from the cortical surface. Measurements usually lasted an hour, but were not acquired here. Time in tissue was 8–30 minutes. The electrode painted with DiD was inserted in the mouse cortex and left for 2 hours. The multichannel silicon electrodes (multi-electrodes) that were used had either 8 shanks, 40 *μ*,m wide, 15 *μ*m thick, 200 *μ*,m apart (Neuronexus inc., part no. A8⨯8-5mm-200-160, where one lateral shank broke off during a previous experiment), 4 shanks, 83 *μ*,m wide, 15 *μ*,m thick, 150 *μ*,m apart (Neuronexus inc., part no. A4⨯1-tet-3mm-150-150-312-A16) or 4 shanks 125 *μ*,m apart (Neuronexus Inc., part no. A4⨯4-3mm-50-125).

### CLARITY

Mice and rats were perfused transcardially with 4% w/v paraformaldehyde (PFA) in phosphate buffered saline (PBS). The spinal cord and brain was then removed and stored in 4% w/v PFA overnight. All tissue was stored in PBS until further processing. Brain tissue was embedded in 3% w/v SeaPlaque^®^ Agarose (Lonza) and cut with a vibratome (VT1000 S, Leica Biosystems) into 1000 *μ*,m slices. Spinal cord segments were manually cut with a scalpel along the midline or embedded in agar as brain tissue and cut into 600 *μ*,m transverse slices. The CLARITY clearing processing was performed via PACT: a passive CLARITY techniqu^e59,60^and accomplished according to Yang et al. 2014^13^ by the following steps: The tissue was placed in a conical tube with 4% w/v acrylamide, 0,05% w/v bis-acrylamide, 0.25% w/v VA-044 (Wako Chemicals) in PBS, and incubated overnight at 5°C. The following day, the solution was degassed with N2 for 5 minutes, then incubated at 37°C while gently shaking for 4 hours until the hydrogel had fully polymerized. The tissue was extracted from the gel with Kimwipes and washed in 8% w/v sodium dodecyl (SDS, Sigma-Aldrich) in PBS, pH 7.5, twice over a day at room temperature or 37°C. The clearing process continued over several nights with replacement of SDS each day if possible to enhance the clearing speed. Coronal mouse brain sections (1 mm thick) and (1 cm long) adult mouse spinal cords segments were cleared over 4 days, while rat spinal cord segments were cleared over 6 days.

### Staining and Mounting

After the CLARITY procedure, the sample was washed in PBS-T (PBS with 0.1% v/v Triton-X-100) at room temperature for 10 minutes during gentle shaking, which was repeated once. Nuclear DNA was stained with 2 *μ*g/ml DAPI in PBS with 0.1% v/v TritonX-100, and 0.01% w/v sodium azide, at room temperature during gentle shaking over two nights. After staining, the tissue was washed twice with PBS for 10 minutes and gently dried with filter paper. Then tissue was transferred to a refractive index matching medium (RIMM) of either 80% v/v glycerol (Sigma-Aldrich Co), 88% w/v Histodenz (Sigma-Aldrich Co), or 63 v/v % 2,2’-thiodiethanol (TDE, VWR international) in PBS with 0.01% sodium azide and pH adjusted to 7.5 with NaOH. The tissue was incubated in RIMM at room temperature until transparent, usually overnight, prior to imaging. The tissue was placed in a chamber designed for live cell imagining (POC R2, PeCon) consisting of two 0.17 mm glass slides with silicon spacers to prevent compression. The tissue was sandwiched between the glass slides while being covered in RIMM and gently held in place without being compressed by the silicon spacers.

### Imaging

We used confocal microscopes LSM 700 and LSM 710 (Zeiss) with EC Plan-Neofluar, air 10x/NA 0.3 with a working distance of 5.2 mm. Excitation sources were a 30 mW diode for 405 nm, a 25 mW Argon laser at 488 nm and 514 nm (CM-DiI and FM 1-43FX were excited at 5% and SP-DiI at 10% laser power) with an Airy unit of 0.98-1.5. Images were acquired with 2 or 4 times frame-averaging, tiles had 0–10% overlap and Z-stacks had 10% overlap. Overview images were acquired with a Zeiss SteREO Lumar V12 with a 1.5× Neolumar S. Post-processing (contrast adjustments, and subtraction in Fig. 3j.-l and Supplementary Fig. 3.e: 50% of the DAPI, GFP and DiIs images), maximum intensity projections (MIP) and 3D-reconstructions) was done in Zen Blue 2012 (Zeiss) and Vaa3D 3.055.^61^ Macroscopic images where taken with a M320 Dental Microscope (Leica).

### Fluorescent measurements

We measured the fluorescence of the electrode traces seen in the confocal images (Fig. 3) on unadjusted MIPs. The mean intensity in 80 *μ*m × 600 *μ*m rectangles placed on the identifiable electrode traces were subtracted the mean background intensity in a 160 *μ*m × 600 *μ*m rectangle placed elsewhere on the tissue in the images. This was done in Zen Blue 2012 (Zeiss). Measurements were handled in Prism 6 (GraphPad).

## Acknowledgements

We thank Peter Petersen for assistance with multielectrode dye-painting and placement. Rasmus Christensen inserted DiD painted electrode, and R. Christensen, Eva Carlsen, Tania Barkat, and Michael Nielsen provided rodents. Thanks to the Core Facility for Integrated Microscopy, Faculty of Health and Medical Sciences, University of Copenhagen, and their assistance provided by Thomas Braunstein and Laura Plantard. This research was supported by the Novo Nordisk Foundation (RB) and The Danish Council for Independent Research Medical Sciences (RB) and The Medical Society in Copenhagen (KJ). The work is part of the Dynamical Systems Interdisciplinary Network, University of Copenhagen.

## Author contributions statement

KJ and RB conceived and conducted the experiment, and analysed the results. Both authors reviewed the manuscript.

## Additional information

The authors declare no conflicts of interest.

